# Cellular and Synaptic Diversity of Layer 2-3 Pyramidal Neurons in Human Individuals

**DOI:** 10.1101/2021.11.08.467668

**Authors:** Henrike Planert, Franz Xaver Mittermaier, Sabine Grosser, Pawel Fidzinski, Ulf Christoph Schneider, Helena Radbruch, Julia Onken, Martin Holtkamp, Dietmar Schmitz, Henrik Alle, Imre Vida, Jörg Rolf Paul Geiger, Yangfan Peng

**Author notes:** These authors contributed equally (Henrike Planert, Franz Xaver Mittermaier, Sabine Grosser). Senior and corresponding authors^‡^,.

## Abstract

Understanding the functional principles of the human brain requires deep insight into the neuronal and network physiology. To what extent such principles of cellular physiology and synaptic interactions are common across different human individuals is unknown. We characterized the physiology of ~1200 pyramidal neurons and ~1400 monosynaptic connections using advanced multineuron patch-clamp recordings in slices from human temporal cortex. To disentangle within and between individual sources of heterogeneity, we recorded up to 100 neurons per single subject. We found that neuronal, but not synaptic physiology varied with laminar depth. Connection probability was ~15% throughout layer 2-3. Synaptic amplitudes exhibited heavy-tailed distributions with an inverse power law relationship to short term plasticity. Neurons could be classified into four functional subtypes. These general principles of microcircuit physiology were common across individuals. Our study advances the understanding of human neuron and synaptic diversity from an individual and phenotypic perspective.

## Introduction

Cellular diversity provides biological tissue with the capability to fulfil a multitude of functions. Recent advances in single-cell technology uncovered great cellular heterogeneity across different human tissues^1^, including that of the human neocortex. Here, high-throughput molecular approaches uncovered transcriptomic neuron types that were broadly conserved across species^2,3^. While superficial neocortical layers are generally assumed to comprise rather homogeneous pyramidal neurons, the human layer 2-3 (L2-3) exhibits an increased diversity of pyramidal neuron subtypes^3,4^, and these correspond to diverging electrophysiological and morphological characteristics^5,6^. While the above-mentioned studies defined pyramidal neuron subtypes in L2-3 based on either molecular profile^3^ or morphology^4^, a functional classification based on physiology is lacking. It would provide a complementary phenotypic perspective on neuronal diversity that also incorporates modifications after transcription and translation^7,8^.

In addition to cellular diversity, network function relies on the synaptic connections between neurons. Previous studies on human synaptic physiology uncovered distinct properties within and across cortical layers^9–15^. A detailed characterization of synaptic diversity within an intralaminar cortical microcircuit is of great scientific interest as both neuronal and synaptic heterogeneity have been suggested to improve network computation and stability^16,17^.

In human studies it is especially challenging to derive general principles considering the differences in genetic and environmental backgrounds and their influence on functional properties of the neurons^18–20^. Previous studies on human neuronal and synaptic function relied on electrophysiological approaches with a limited number of samples from each individual ^4,5,9,10,12,14,18,20–23^, requiring pooling across many different individuals. Thus, generalization of cortical microcircuit properties is still limited by the unresolved contributions of within- and between-individual differences to functional heterogeneity.

In order to identify organizational principles of the human cortical L2-3 microcircuit across individuals, we characterized cellular and synaptic properties from large samples of pyramidal neurons per single subject. We performed multineuron patch-clamp recordings on human brain slices from resected temporal cortex of epilepsy patients, followed by anatomical reconstruction^11^. By investigating the human L2-3 cortex in unprecedented detail, we identified principles of within-individual pyramidal neuron and synaptic diversity.

## Results

### Human multineuron patch-clamp recordings with large data samples per individual

We performed multineuron patch-clamp recordings in acute brain slices from the human temporal cortex (see Methods). The cortical tissue was obtained from 23 patients that were diagnosed with pharmaco-resistant epilepsy and underwent anterior temporal lobe resection surgery (Figure 1A). To allow for generalization of findings at the level of individuals, our study was designed to maximize experimental yield per tissue donor^11^ (see Methods). For this study, we focused on cortical subregion L2-3, recording from neurons down to 1200 μm laminar depth from the pia. Studying L2-3 of the human brain is of particular interest since these layers expanded disproportionally during human evolution^24^. Up to ten neurons were recorded simultaneously in whole-cell configuration, allowing us to functionally characterize these cells and to probe for synaptic connections between them (Figure 1A-C). Recorded neurons were filled with biocytin and subsequently visualized (Figure 1B). Out of 1479 pyramidal neurons recorded across all individuals, 1214 met our quality control criteria for inclusion (see Methods, Figure S1). 901 neurons from 22 individuals met further requirements (complete data for functional cellular properties and coordinates, resting membrane potential between −90 and −60 mV) and were used for cellular property analyses (Figure 1F, see Methods, Figure S1). Across all individuals we tested a total of 9834 potential connections between pyramidal neurons and detected 1419 monosynaptic connections by evoking excitatory postsynaptic potentials (EPSPs) in response to elicited presynaptic action potentials (APs, see Methods, Figure 1C-E). On average, we recorded 53 pyramidal neurons and 62 monosynaptic connections per patient (Figure 1F). In summary, we acquired an extensive dataset of cellular and synaptic physiology from the human L2-3 pyramidal neuron population with the potential to address within-individual microcircuit principles.

**Figure 1:**
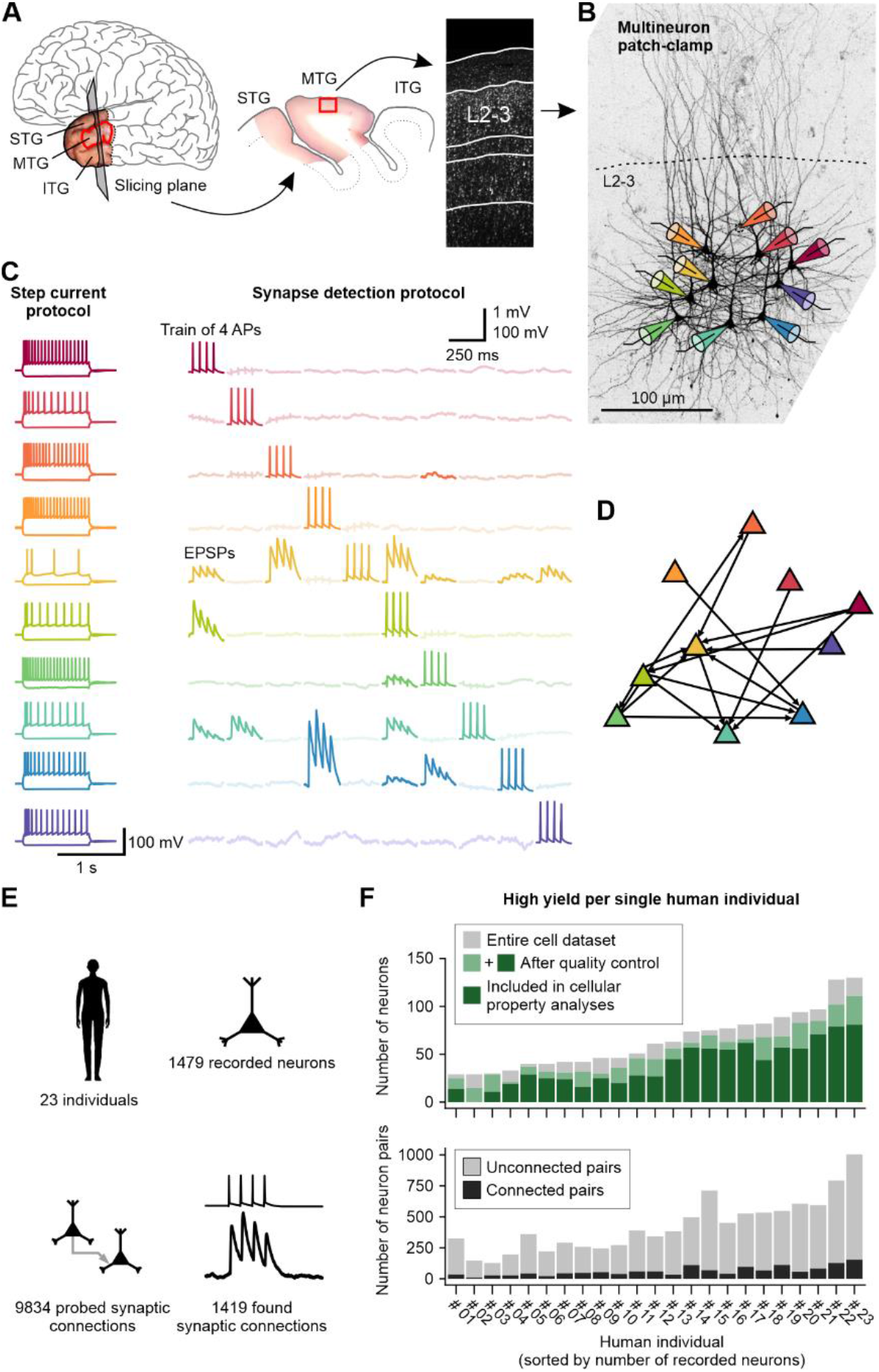
Multineuron patch-clamp recordings of cortical layer 2-3 pyramidal neurons with high yield per individual. (**A**) Human cortical brain tissue was obtained from patients that underwent anterior temporal lobe resection for treatment of epilepsy. Left: Colored photograph of resected temporal lobe tissue mapped onto the human brain. It comprises the anterior parts of the superior, medial and inferior temporal gyrus (STG, MTG, ITG). This study mostly focused on acute slices from the MTG that were cut along the coronal slicing plane. Middle: Photograph of an acute brain slice. Red square marks the cortical region of interest. Recordings were made in L2-3 as depicted in the post-hoc Nissl staining on the right. (**B**) Biocytin staining of recorded neighboring pyramidal neurons. Colored patch pipettes illustrate the multineuron patch-clamp approach (dashed circle: one neuron was not stained). (**C**) Electrophysiological traces recorded from the 10 pyramidal neurons depicted in B. Each row corresponds to one neuron. Left column shows action potential firing upon step depolarization. Right matrix of traces shows averaged postsynaptic voltage traces during subsequent stimulation of action potentials in the neurons (diagonal). Correlated excitatory postsynaptic potentials are highlighted and indicate a monosynaptic connection onto this neuron. Vertical scale bar: APs 100 mV, EPSPs 1 mV. (**D**) Connection pattern between the recorded neurons based on connectivity screening. (**E**) Summary statistic of the whole dataset. The multineuron patch-clamp experiments were repeated multiple times per individual and data was pooled across 23 individuals. (**F**) The number of recorded neurons and number of connected and unconnected neuron pairs are shown for each sampled human individual. Individuals are sorted by the number of recorded neurons and numbering is retained throughout the figures.

### Human pyramidal neuron function changes with laminar depth

Phenotypic differences between pyramidal neurons have been mostly attributed to subtypes between cortical layers. Previous studies of human pyramidal neuron physiology have shown that specific electrophysiological properties, such as the input resistance, sag ratio and AP kinetics, depend on the laminar depth of pyramidal neurons^5,9,21,25^. To assess determinants of neuronal diversity in our dataset, we functionally characterized human L2-3 pyramidal neurons. We extracted 15 passive and active electrophysiological properties that are frequently used to assess the functional phenotype of neurons^4,5,21,25–27^ (Figure 2A, see Methods). These properties showed broad and, in some cases, skewed distributions (Figure 2B). We used AP halfwidth as an example property to demonstrate that distributions of individual subjects mostly resembled the pooled distribution of a respective parameter (Figure 2C). Accordingly, subject medians for pyramidal cell properties were rather similar while there was large variance in each subject. In accordance with previous studies, we found significant relationships between laminar depth and input resistance (R^2^ = 0.06; F-test, p < 0.0001), F/I slope (R^2^ = 0.04; p < 0.0001), sag ratio (R^2^ = 0.04, p < 0.0001) and additional cellular properties (Figure 2D-F). We further found depth dependency of input resistance within individual human subjects (Figure 2G) and used a linear-mixed effects model with varying slopes for subjects to predict that across all individuals the slope is –69.9 ± 56.6 MΩ/mm (fixed slope ± s.d.; Figure 2H, see Methods). The depth dependency could be generalized across individuals for other properties as well, such as F/I slope and sag ratio (Figure S2). This confirms that laminar depth-dependent change of neuronal function is a ubiquitous principle in the human temporal L2-3 cortex.

**Figure 2:**
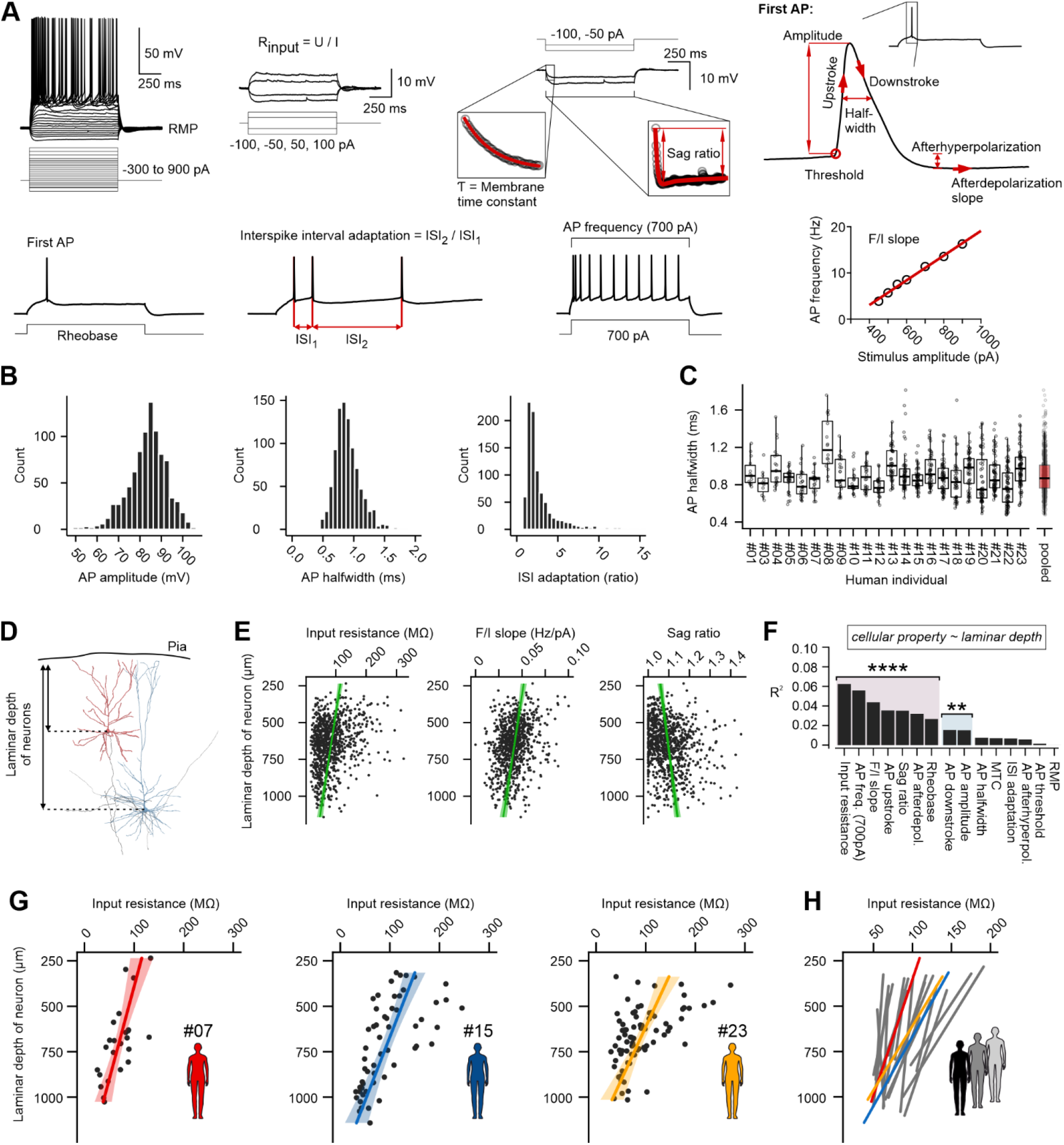
Functional properties of L2-3 pyramidal neurons. (**A**) Schematics depicting the electrophysiological parameters extracted from the raw traces by an automated analysis pipeline. For this study 15 well established electrophysiological properties were analysed (see Methods). (**B**) Histograms show distributions of exemplary electrophysiological parameters. (**C**) Jitter- and boxplots of AP halfwidth for each of the human subjects and for data pooled across all subjects (**D**) Schematic showing how laminar depth of a neuron was measured. (**E**) Scatterplots show input resistance, F/I slope and sag ratio in relationship to laminar depth for the pooled dataset. Green lines and shaded area correspond to regression lines and 95% confidence intervals (c.i.). (**F**) Bar graph shows R^2^ values for each of the 15 cellular parameters obtained from linear regression models (F-test, Bonferroni-corrected; adjusted p-values are indicated by asterisks, ** < 0.01, **** < 0.0001, asterisks apply to all parameters underneath the line). (**G**) Relationship between input resistance and laminar depth in three exemplary human subjects (colored lines and shaded area correspond to regression lines and 95% c.i.). (**H**) Fit lines obtained from linear mixed-effects model with varying slopes and intercepts for each human subject (see Methods). Abbrev.: RMP, resting membrane potential, MTC, membrane time constant, ISI, inter-spike interval.

### Principles of connectivity and synaptic properties of human L2-3 pyramidal neurons

Microcircuit functions do not only rely on intrinsic cellular properties but also on synaptic interactions. We used our multineuron patch-clamp approach to investigate monosynaptic connections between simultaneously recorded neurons in human L2-3. First, we assessed connection probability, which is the ratio of synaptically connected neuron pairs to all probed pairs. We excluded slices with a slicing angle non-parallel to the orientation of apical dendrites and neurons with cut axons to avoid underestimation of synaptic connectivity (see Methods). Mean connection probability between L2-3 pyramidal neurons in the temporal cortex was 15.8% (1094/6911), in line with previous studies^9–12,14^. At the level of individual humans, the connection probability ranged from 6.8% to 24.0% (Figure 3A), differing significantly across individuals (χ^2^ test, p < 0.0001). In our dataset, recorded neuron pairs had intersomatic distances of 128 μm ± 55 (mean ± s.d.) and laminar depth varied between 242 and 1122 μm (Figure 3B-C). As expected, connection probability decreased with increasing intersomatic distance (Figure 3C)^9,28^.

**Figure 3:**
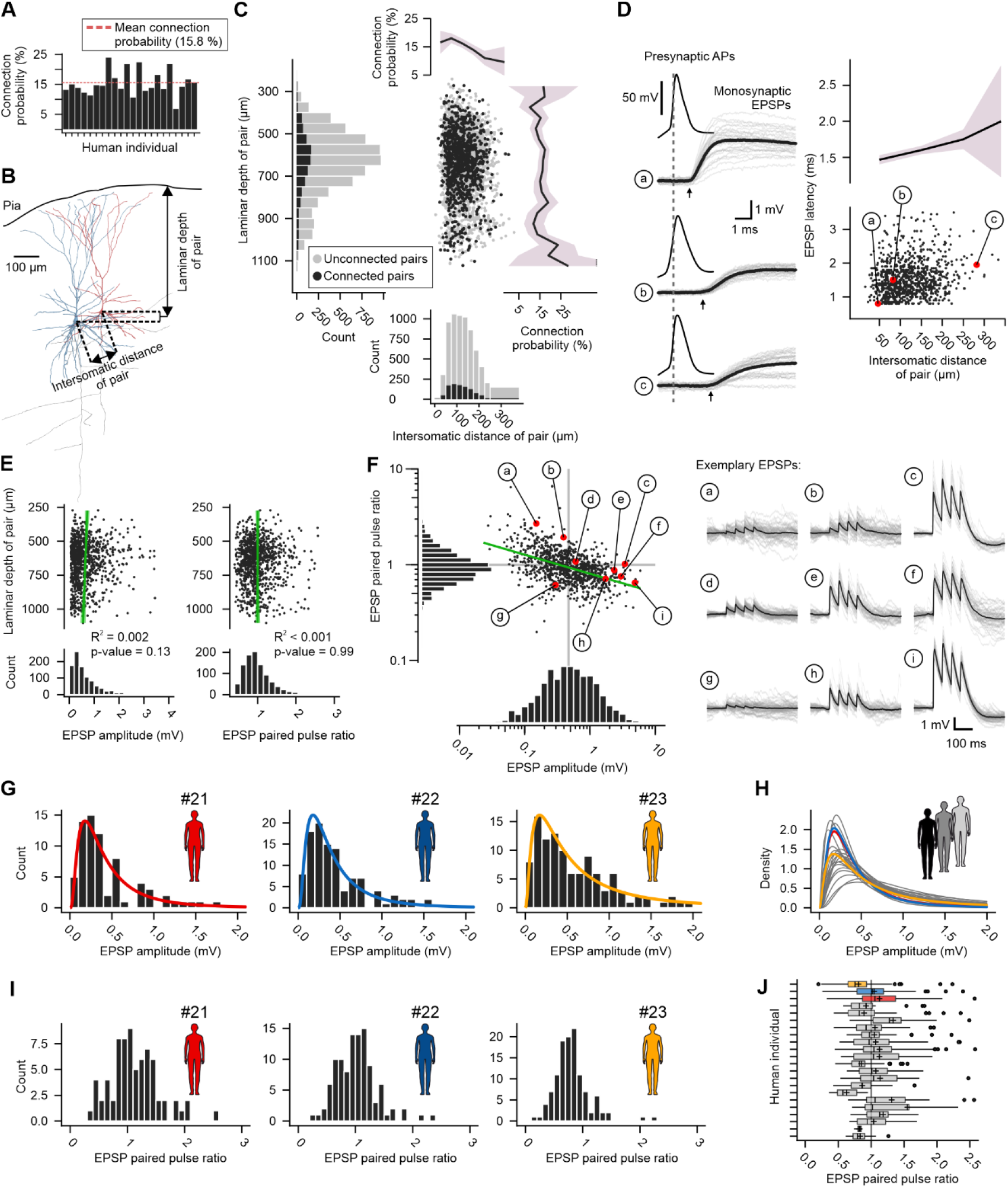
Connectivity and properties of synaptic connections between human L2-3 pyramidal neurons. (**A**) Bar graph depicting pyramidal neuron connection probability for each individual. Dashed red line represents the mean connection probability across all individuals. (**B**) Schematic demonstrating measurement of laminar depth and intersomatic distance of a neuron pair. (**C**) Scatterplot and histograms show the intersomatic distance and laminar depth distribution of all recorded neuron pairs. Black dots represent pairs that are connected by a synapse, grey dots are tested but unconnected pairs. Line plots on top and on the right show distance- and depth-dependent connection probabilities (line represents mean connection probability for bins; shaded area represents bootstrapped 95% c.i.). (**D**) Traces recorded from pre- and postsynaptic neurons of three example synapses (a, b, c) are plotted on the left. Selected synapses display different latencies from presynaptic AP to the EPSP onset (marked by arrows). Scatterplot on the right shows increasing synaptic latency in relation to the intersomatic distance of the pre- and postsynaptic neuron. Black line and shaded area on top represent the mean and 95% c.i. of 100 μm bins. (**E**) Scatterplots show EPSP amplitude and paired pulse ratio (PPR) in relationship to laminar depth (green lines and shaded area correspond to regression lines and 95% c.i.). Histograms on the bottom show skewed distributions (x-axes are cropped for better visualization, see F for all data points). (**F**) Log-log plot shows power law relationship (green line) between EPSP amplitude and PPR. Note that histograms of logarithmically displayed data show symmetrical distributions. Vertical and horizontal grey lines correspond to the median EPSP amplitude and PPR of 1.0, respectively. Red dots labelled with lowercase letters mark example EPSPs that are plotted on the right (black trace represents average from single sweeps in grey). (**G**) Histograms show right-skewed distribution of EPSP amplitude for three exemplary human subjects. Colored lines reflect log-normal fits of the data. (**H**) Overlaid log-normal fit lines for each of the individuals. (**I**) Histograms show distribution of EPSP PPR for three exemplary human subjects. (**J**) Boxplot shows distribution of PPR for all individuals.

In addition to connectivity, the computational capacity of a cortical microcircuit is constrained by the phenotypic diversity of its synapses^16^. We measured synaptic properties from averaged EPSP traces, such as synaptic latency, maximum slope, amplitude, halfwidth and paired pulse ratio (PPR) as a proxy for short-term plasticity (see Methods). Monosynaptic EPSPs exhibited onset latencies of 1.56 ± 0.65 ms (mean ± s.d.) relative to presynaptic somatic AP. This synaptic latency consists of different delays, such as the axonal, synaptic as well as postsynaptic dendritic delay. The axonal delay depends on the axonal conductance speed, which was recently reported to be in the range of 1.5 m/s in human neurons^29^. In our data, the synaptic latency showed a positive and nearly linear relationship with intersomatic distance (1.7 ms/mm; F-test, p < 0.0001, Figure 3D). Many synaptic connections had small EPSP amplitudes (< 1 mV), and only very few had large amplitudes (>3 mV; Figure 3E-F). This skewed distribution of synaptic amplitudes has been observed across cortical layers and areas^30–33^, and large-amplitude synapses brought about by plasticity may underlie the coactivation of groups of neurons thought to be fundamental for computations of the brain^34–36^. The median amplitude of 0.46 mV is within the range of previous human studies^9,10,12,14,37^. Similar to EPSP amplitude, the PPR showed a skewed distribution (Figure 3E-F). The median of 0.93 is also in overall accordance with previous short-term plasticity measures at 20 Hz stimulation^9,37^. Given the apparent log-normal distribution of both synaptic amplitude and PPR and their expected inverse correlation, we analysed their relationship in a log-log plot. We could show that these two parameters follow a power law relationship with an exponent of −0.2 and a coefficient of 0.8 at 20 Hz stimulation frequency (Figure 3F). To validate that the large heterogeneity of synaptic strength and dynamics is a general principle within each individual and not a result of pooling across diverse human subjects, we analysed the properties at the level of individual human subjects. We fit the EPSP amplitude distribution of each subject with a gaussian and a log-normal distribution and found that the log-normal fits were a better model of the data in all individuals (Figure 3G-H, see Methods). This suggests that a skewed distribution of amplitudes is a ubiquitous feature within each human individual, highlighting it as a fundamental principle of synaptic diversity^38^. For the EPSP PPR we were able to show that all but two human subjects had both paired pulse depressing (PPR < 1.0) and paired pulse facilitating synapses (PPR > 1.0) at 20 Hz stimulation (Figure 3I-J).

Given the depth-dependence of cellular properties, we also investigated whether synaptic connectivity and properties correlate with laminar depth. Unlike previously reported^3^, short-term plasticity showed no significant relationship with laminar depth (R^2^ < 0.001; F-test, p = 0.99; Figure 3E). Differences in measures used to assess short term plasticity might explain this diverging result. Similarly, neither connection probability (stable at ~15%) nor EPSP amplitude (R^2^ = 0.002; F-test, p = 0.13) were correlated with laminar depth (Figure 3C). Taken together, synaptic properties exhibit a great diversity within human individuals and, in contrast to some cellular properties, do not change along the laminar depth of L2-3.

### Human L2-3 pyramidal neurons can be classified into functional subtypes

Large diversity of human pyramidal neurons in L2-3, including several subtypes, has been proposed based on transcriptomic profile^3^ and morphology^4^. We investigated this diversity from the phenotypic perspective by exploring the parameter space of 15 frequently used cellular electrophysiological properties. Some of these properties were significantly correlated (Figure 4A). To quantify the overall degree of correlation between these parameters, we conducted a principal component analysis (PCA, Figure 4B). The first principal component explained 24% of the total variance in the cellular dataset, and 7 out of 15 principal components were necessary to capture ~80% of the variance. Thus, while some degree of correlation was identified, the functional parameter space of pyramidal neurons appears to be rather high-dimensional. Furthermore, we found that the within-individual variance was severalfold higher than the between-individual variance across cellular properties (Figure S3). This suggests that pyramidal neurons within the temporal cortex of a single human can exhibit strongly divergent properties (Figure S3B). To quantify this observation, we calculated the intraclass correlation coefficients (ICC) from random-effects models (see Methods). The ICC can be interpreted as the fraction of the total variance of a parameter that is explained by a certain grouping structure in the data, in our case individuals^39^. We found that differences between individuals only accounted for 3.7 - 29.7% of the total variance of cellular properties (Figure S3C). The results of the PCA and the random-effects models suggest that variability of cellular electrophysiological properties of pyramidal neurons is high-dimensional and can largely be attributed to within-individual cellular diversity.

**Figure 4:**
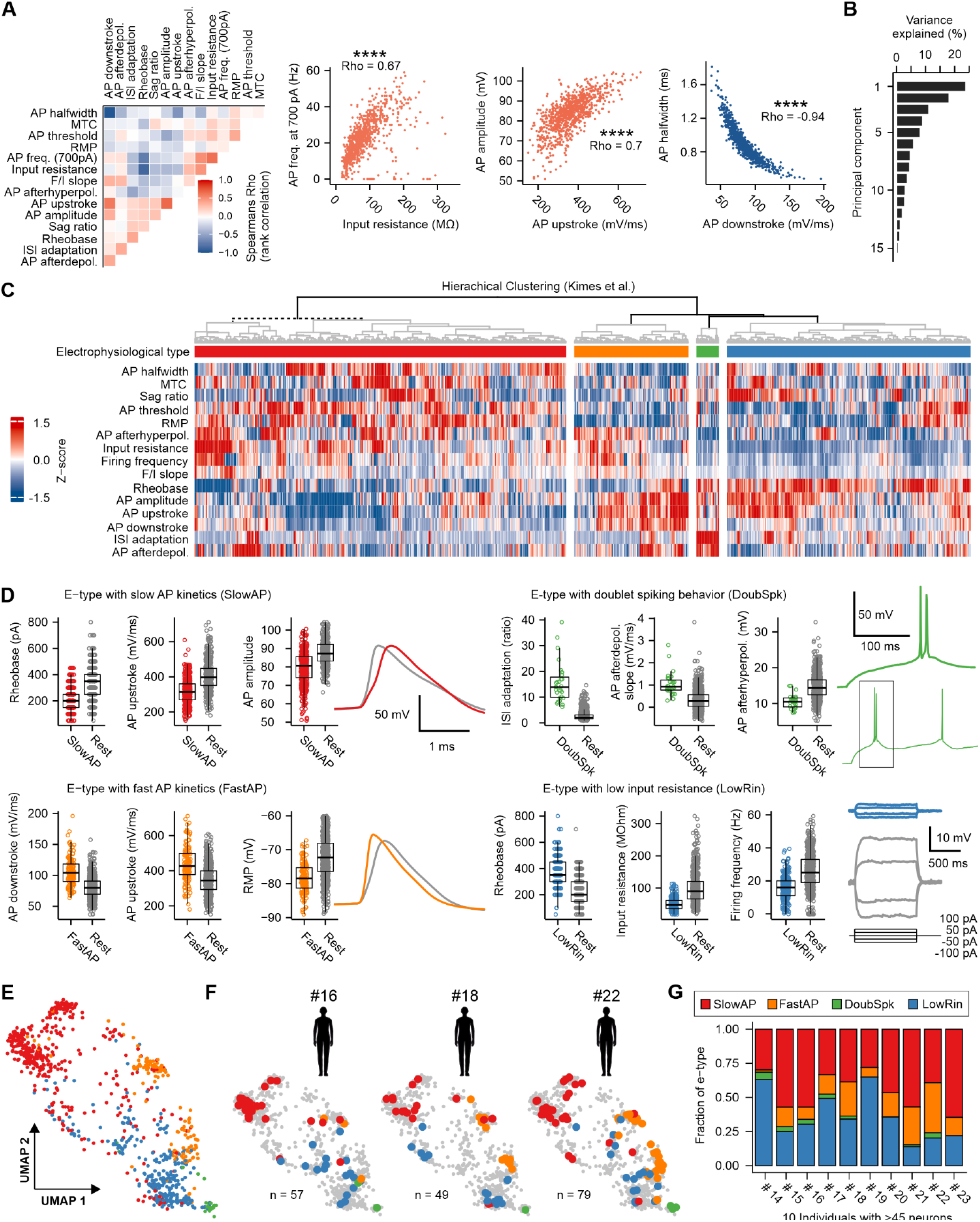
Clustering of human L2-3 pyramidal neurons into four electrophysiological types. (**A**) Left: Matrix shows correlation coefficients for each pair of parameters. Right: Parameter pairs with strong correlation are shown as scatter plots with the color of data points corresponding to the correlation coefficient of the respective parameter pair (Spearman rank correlation test, p-values are indicated by asterisks, **** < 0.0001). (**B**) Scree plot of a principal component analysis of the 15 electrophysiological properties shows the fraction of variance explained by each principal component. (**C**) Clustergram visualization of the cellular electrophysiological parameters for all pyramidal neurons. Each column represents one neuron, and the color code represents the Z-score of each parameter (rows). Neurons are grouped into four electrophysiological types (e-types) according to hierarchical clustering using Euclidean distance and Ward’s linkage criterion. The color of the dendrogram on top indicates the result of statistical testing of the bifurcations (see Methods): Black indicates significant bifurcations (p-values < 0.05, corrected for family wise error), dashed line indicates non-significant bifurcation, grey indicates untested bifurcations due to cluster size below threshold. (**D**) The three most distinguishing parameters of each e-type are shown. Boxplots display the comparison between the neurons of a respective e-type (color code) with all other neurons (grey) for a given parameter. Example traces are shown to the right of the respective boxplots (colored trace corresponds to an exemplary neuron of the respective e-type, grey trace corresponds to a neuron which belongs to a different e-type). (**E**) Uniform Manifold Approximation and Projection (UMAP) visualization of the high-dimensional parameter space. Each dot represents one neuron color-coded by assigned e-type. (**F**) Three separate UMAPs showing neurons recorded from selected individuals with large sample sizes. Note that all four e-types are present in each of the example individuals. (**G**) Stacked bar plot shows relative fraction of e-types for ten patients, for which more than 45 pyramidal neurons were recorded.

Given the large within-individual phenotypic diversity of pyramidal neurons, we asked whether they occupy a continuum in the high-dimensional functional parameter space or, alternatively, are grouped into distinct subspaces possibly reflecting functional cell types^8,40^. To address this, we performed an unsupervised hierarchical cluster analysis based on the 15 electrophysiological properties^41,42^. The cluster algorithm was followed by a Monte Carlo approach to test for statistical significance of cluster separation (Figure 4C)^43^. We further performed a resampling analysis to find an appropriate number of clusters and establish robustness of the clustering (see Methods, Figure S4)^44^. Based on our set of parameters, these analyses allowed us to propose four electrophysiologically distinct subtypes (e-types) of pyramidal neurons in L2-3 (Figure 4C-D, Figure S4G). To test whether other clustering algorithms yield similar results we performed k-means clustering into k=4 clusters with randomly selected starting points (see Methods). The k-means algorithm separated the pyramidal neurons into four very similar groups compared to the hierarchical clustering (Figure S4H). To better understand the e-type clustering result, we identified the most distinguishing features for each e-type (Figure 4D, see Methods). Over one third of neurons belonged to an e-type that was characterized by a low rheobase, comparatively slow AP upstroke as well as low AP amplitudes and was thus termed “SlowAP”. In contrast, the “FastAP” e-type exhibited faster AP up- and downstroke kinetics and more negative resting membrane potentials. The “DoubSpk” e-type represented a small group of neurons with initial doublet spiking, characterized by a large inter-spike interval adaptation, steep afterdepolarization slope and low AP afterhyperpolarization. Finally, the “LowRin” e-type was overall less excitable, characterized by a high rheobase, low input resistance and lower AP frequency at 700 pA stimulation. To visualize the e-type clustering in the high-dimensional parameter space, we performed dimensionality reduction^45^ (see Methods), which revealed cell clusters with broad distributions and some regions of overlap (Figure 4E). In order to determine whether the presence of these e-types represents a general principle across individuals, we compared the clustering result of pyramidal neurons at the level of single individuals. UMAPs showed similar patterns of distribution across individuals (Figure 4F). Additionally, we assessed the fraction of e-types in the ten individuals with more than 45 sampled neurons and found that the three most common e-types were present in each of these individuals. Only the small group of “DoubSpk” e-type was absent in the data of three humans, possibly reflecting a sampling limitation (Figure 4G). While the relative proportions of e-types differed across individuals, they largely resembled the overall fractions. Thus, our analysis of the high-dimensional cellular parameter space showed largely preserved e-types across individuals. This supports a functionally relevant subdivision of pyramidal neurons and adds to the evidence of increased pyramidal neuron diversity as a general principle in human cortical L2-3^3–5,21,25^.

## Discussion

Physiology of neurons and their synapses in the human neocortex has been challenging to study due to sample size limitations of electrophysiological approaches and potential high variation between individuals, such as differences in genetic background and life history. In this study, we took advantage of a high-throughput multineuron patch-clamp approach^11^ to study cellular and synaptic properties at the level of single subjects. We identified principles of functional and synaptic diversity of L2-3 pyramidal neurons that were preserved across individual humans.

By recording over 1200 pyramidal neurons in human L2-3 temporal cortex, we were able to propose four electrophysiological subtypes. Previous studies found a higher transcriptomic diversity of L2-3 pyramidal neurons in human compared to mouse cortex, suggesting five t-types in the supragranular pyramidal neuron population^3,5,46^. Two of these t-types were restricted to deep L3, which is likely outside the range sampled in our study (Figure 5B)^5^. In addition to gene expression, the functional classification in this study incorporates changes in physiology due to posttranscriptional regulation, plasticity, and neuromodulation^8,47^. In line with this, enrichment of many proteins in the brain was found to be in discordance with the transcriptome expression^48^. Thus, whether the e-types identified here are transcriptomically distinct or represent functional states or gradients within or across t-types remains to be determined^8^. Distinguishing between cell types and functional states with respect to different levels of granularity remains an ongoing challenge and is the focus of large-scale single-cell transcriptomic efforts^8,14^. Here we provide a comprehensive characterization from the phenotypic perspective.

We analysed over 1000 monosynaptic connections between pyramidal neurons and were able to corroborate known and identify novel principles of human synaptic physiology. We found overall connection probability between L2-3 pyramidal neurons to be 15.8% with between-individual variations. This is slightly higher compared to the ~13% which we calculated by pooling across previous human multipatch studies that reported in total ~200 synapses between L2-3 pyramidal neurons^9,10,12,14,37^. Our data were able to capture the skewed distribution of synaptic amplitudes with many weak and few strong connections across laminar depth and in each individual human. As this feature has been ubiquitously reported in excitatory and inhibitory synapses across brain regions and species^9,12,31,38,49^, our within-individual result further establishes it as an evolutionary conserved principle of synaptic transmission that is likely important for network computation and plasticity^30,38^. We further found paired pulse ratio of amplitudes elicited at 20 Hz to exhibit a log-log relationship with synaptic amplitude. While previous studies reported paired pulse depression at higher stimulation frequencies^9,37^, our results suggest that human excitatory synapses can exhibit both depression and facilitation at an intermediate stimulation frequency. In contrast to some cellular properties, neither of these synaptic parameters varied with laminar depth up to 1.2 mm. Overall, our large synaptic dataset obtained in homogenous experimental conditions captured highly robust and general principles of human synaptic properties within individuals.

Differences in single neuron properties between individuals have been suggested to correlate with intelligence and have been hypothesized to underlie differential network performance^5,15,21,22,50,51^.

Interindividual differences of neuronal physiology have also been described in animals^52^. While single parameters likely affect network performance, it remains to be shown whether interindividual differences of these parameters could represent distinct solutions to achieve similar computations^53^. Our data suggest that individuals are surprisingly similar in average cellular and synaptic properties with extensive within-individual diversity. Such diversity of cellular and synaptic properties within an individual network has been implicated in improved performance and stability^18,54,55^. We thus propose a framework in which diversity of cellular and synaptic components within individuals represent key determinants of human cortical microcircuit functions.

Like all neurophysiology studies on acute human brain slices^5,15,21,22,50,51^, we used resected tissue from neurosurgical patients and focused on cortical tissue that was remote to the pathology. While we cannot completely rule out effects associated with epilepsy or its underlying etiologies, we excluded structural abnormalities through neuropathological assessment of the tissue used. We found no evidence for pathology-related effects on the pyramidal neuron physiology. This is in line with previous studies that have shown an independence of neuronal and synaptic properties from disease state of the patient^56,57^. The fact that we found preserved principles of cellular diversity and synaptic connectivity across individuals despite substantial differences with regard to sex, age, and patient phenotypes is remarkable and supports the generalizability of our study.

## Acknowledgements

We thank Christian Madry, Gilad Silberberg and Jeffrey Stedehouder for comments on earlier versions of this manuscript. We thank Andrea Wilke for technical support.

This research was supported by the Deutsche Forschungsgemeinschaft (DFG): Walter Benjamin Fellowship to YP (451242556), NeuroCure Cluster of Excellence to DS (EXC2049).

## Author Contributions

Initial concept: JG

Conceptualization: HP, YP, JG, FM, HA, SG, IV

Project management: YP

Supervision: JG, YP, IV

Patient material: PF, US, JO, MH

Electrophysiological data acquisition: HP, FM, YP

Anatomical data acquisition: SG

Neuropathological assessment: HR

Data software development: FM, YP

Cellular physiology analysis: FM

Synaptic physiology analysis: YP, FM

Anatomical data analysis: SG

Interindividual analysis: HP, FM, YP

Manual data curation: HA, FM, YP, HP, JG

Visualization: FM

Funding acquisition: DS

Resources: JG, IV, DS

Writing - original draft: HP, YP, FM, HA, SG, IV, JG

Writing - review and editing: all authors

## Methods

### Experimental procedures

#### Human subjects

The data obtained in this study was measured in acute brain slices obtained from temporal lobe resections in 23 patients suffering from pharmaco-resistant epilepsy (12 male, 11 female, age range 21 to 55 years with median age of 34 years). The study procedures adhered to ethical requirements and were approved by the local ethical committee with Approval Nr. EA2/111/14. Prior written informed consent for the scientific use of resected tissue was given by the patients.

#### Human brain slice preparation

Temporal lobe pole tissue resected from the patients was transferred from the operating theatre to the laboratory in sterile and cooled sucrose-based aCSF (in mM: 87 NaCl, 2.5 KCl, 3 MgCl_2_, 0.5 CaCl_2_, 10 glucose, 75 sucrose, 1.25 NaH_2_PO_4_, and 25 NaHCO_3_, 310 mOsm) enriched with carbogen (95% O_2_, 5% CO_2_) within 15-40 min after resection. The pia mater was removed from the surface and the tissue was cut under sterile conditions in ice-cold sucrose-based aCSF to 300 or 400 μm-thick slices. After a 30 minutes recovery period at 34°C, slices were held submerged at room temperature in sucrose-based aCSF. In a subset of experiments, an antibiotic (Minocycline, 2 μM) was added to the incubation solution. Recordings were made up to 52 hours after slicing (interquartile range: 7.5 - 26 hours).

#### Multineuron patch-clamp recordings

Slices were transferred to the recording chambers of two patch-clamp setups optimized for high-throughput microcircuit assessment of rare tissue samples, including automated pipette pressure and pipette cleaning systems^11^. The eight-manipulator setup included the following equipment: Eight PatchStar manipulators (Scientifica), BX-51 WI microscope (Olympus), Orca-Flash 4.0 LT Camera (Hamamatsu), MultiClamp 700B amplifiers (Molecular Devices), Digidata 1550 digitizer (Molecular Devices). The ten-manipulator setup included ten u-Mp micromanipulators (Sensapex), Eclipse FN1 microscope (Nikon), Orca-Flash 4.0 LT Camera (Hamamatsu), MultiClamp700B amplifier (Molecular Devices), Power1401-3A digitizer (Cambridge Electronic Design). Data were low-pass filtered with Bessel filter at 6 kHz and digitized at a sampling rate of 20 kHz (with a small subset of data sampled at 10 kHz). Data acquisition was performed using pClamp 10 (Molecular Devices) or Signal 6 acquisition software (Cambridge Electronic Design).

Patch pipettes were pulled from borosilicate glass capillaries (2 mm outer/1 mm inner diameter; Hilgenberg) on a horizontal puller (P-97, Sutter Instrument Company). They were filled with K-gluconate based intracellular solution containing (in mM) 130 K-gluconate, 2 MgCl_2_, 0.2 EGTA, 10 Na_2_-phosphocreatine, 2 Na_2_ATP, 0.5 Na_2_GTP, 10 HEPES buffer and 0.1% biocytin (290-295 mOsm, pH adjusted to 7.2 with KOH) and had 5-8 MΩ resistance. The liquid junction potential was calculated to be 16 mV and was not corrected for. Neurons were visualized using the differential interference contrast infrared microscope and pyramidal neuron-like soma were selected. To target L2-3, we only included neurons that were recorded at a cortical depth up to 1200 μm (distance from pia). To allow within- and between individual comparisons, we only included data from individuals in which we recorded more than 25 neurons. Whole-cell patch-clamp recordings of these L2-3 neurons were performed at 34 °C under submerged conditions, with continuous perfusion of aCSF containing (in mM) 125 NaCl, 2.5 KCl, 1 MgCl_2_, 2 CaCl_2_, 10 glucose, 1.25 NaH_2_PO_4_, and 25 NaHCO_3_ (300 mOsm). Pipette access resistance and capacitance were compensated throughout the experiment, and remaining effects of these were addressed offline by means of reverse RC filtering (see quality control and defiltering procedure below). In some cases of recording failure, we repatched neurons with a cleaned pipette using our automated “clean2complete” approach^11^. For the assessment of cellular and synaptic electrophysiological properties, we applied a range of stimulation protocols in voltage and current clamp, mostly within 30 minutes of recording (see below).

#### Morphological reconstruction

After recording, slices were fixed overnight in a fixative solution containing 4% paraformaldehyde with 0.1 M Phosphate buffer (PB, pH 7.4) at 4°C for 24 - 48 hours. The biocytin-filled neurons were subsequently visualized using avidin conjugated fluorochrome (Alexa Fluor-647, 1:1000, Invitrogen, UK) in a PB-buffered saline (PBS, 0.9% NaCl) solution containing 3% NGS, 0.1% TritonX-100 and 0.05% NaN3 for 24 hours at 4°C. Slices were rinsed in PBS and then desalted in PB before being mounted in a aqueous medium (Fluoromount-G, Southern Biotech) between cover slips with a 300 μm thick metal spacer to prevent shringkage (Bolduan *et. al*, 2020). Visualized neurons were imaged on a laser scanning confocal microscope (FV1000, Olympus, Japan) for morphological identification. Selected neurons were reconstructed using the Neutube software package (Feng et al. 2015) from composite image-stacks obtained with a 30x silicone immersion objective (N.A. 1.05, UPlanSApo, Olympus) over the full extent of the cells.

#### Neuropathological assessment

To confirm cortical tissue quality, parts of the temporal lobe pole tissue, including slices from brain slice preparation, were selected for histopathological evaluation by a board-certified neuropathologist. The analyzed slice samples revealed no structural changes. A subset of tissue samples of two patients exhibited epilepsy-associated pathology (low grade glioneuronal tumor and focal cortical dysplasia), or microinfarcts. All these pathologies were in the temporal lobe but not in the examined tissue used for slices. Patch-clamp recordings of these three patients also displayed no obvious differences in cellular electrophysiology. This, together with the observation that the ~75% anatomically analysed neurons of the dataset exhibited typical pyramidal neuron morphology (see below), suggests that the recorded pyramidal neuron properties were not affected by pathology.

### Cellular physiology analysis

We developed custom MATLAB code to perform automated analysis of recorded raw traces, comparable to previous studies^12,27^, as outlined below.

#### Resting membrane potential, input resistance, membrane time constant and sag ratio

To estimate the *resting membrane potential*, we took average amplitudes of four 150 ms trace segments, located before the −50, 50, 100 and 150 pA step current injections. The mean of the four values was used as the final *resting membrane potential*. The *input resistance* was calculated as the difference between the steady state voltage and the *resting membrane potential*, divided by the injected current, using −100, −50, 50, 100 pA current steps. The average voltage of the last 100 ms before the end of the step current stimulus was defined as the steady state^27^. The mean of the *input resistance* values at the different step current injections was used as the final *input resistance*. To estimate the *membrane time constant*, the *τ*-values of mono-exponential functions (1), which were fit to the initial segments of the voltage responses to the −50 and −100 pA step currents, were utilized. We used the mean of the two *τ* values as the *membrane time constant*.

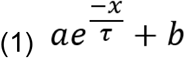

To approximate the sag, we fit the voltage response to a −100 pA step current with a function of the form (2). The sag ratio was calculated from the model fit as the ratio of the negative peak voltage amplitude measured from resting membrane potential and the respective amplitude of the steady state voltage during current injection.

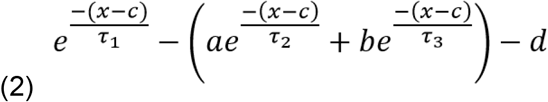

#### Defiltering of action potential traces

APs are fast signals and thus prone to filtering artifacts. Since it is not possible to compensate the entire capacitance of the patch-clamp pipette in current clamp mode, its remainder (referred to as parasitic capacitance^58^), combined with the access resistance, form a filter. Some AP parameters extracted from the original traces show a correlation with the access resistance due to an increasing effect of this filter with larger access resistance (Figure S5). The AP upstroke and the AP amplitude are affected the most (Pearson correlation coefficient is −0.6 for both correlations). Even if one sets strict limits for the access resistance of included recordings, this filter artifact will still affect AP shape parameters recorded with low access resistance. Since we measured the access resistance, and can estimate the parasitic capacitance, we can hence approximate the effect of the filtering. In an approach similar to that of Jayant et al.^59^ we used an inverse digital RC-filter algorithm to produce a defiltered AP signal for each neuron (Figure S5A). The parameters extracted from the defiltered signals show a much weaker correlation with the access resistance (Figure S5B-C). Correcting for the dependence of parameters on the access resistance allowed further analysis with minimized technical bias.

#### Action potential parameters

AP parameters were extracted from the first AP elicited in response to a series of increasing step current injections (increments of 50 pA). All measurements were taken from the defiltered trace. The *AP threshold amplitude* was measured on the ascending trace where the slope exceeded 20 mV/ms. The maximum and the minimum of the AP slope were defined as the *AP upstroke* and *downstroke*,respectively. The *AP amplitude* was measured from the *AP threshold* to the maximum peak. The *AP halfwidth* is the width of the AP (in milliseconds) at half AP *amplitude*. The *AP afterhyperpolarization* was measured as the difference between the *AP threshold* and the minimum after the AP. If no minimum was reached in a 10.0 ms time window, the amplitude at the point after the AP, where the slope exceeded −0.2 mV/ms was used instead. The *AP afterdepolarization slope* was defined as the mean slope of the voltage trace in a 1.0 ms time window after the *AP afterhyperpolarization*.

#### Firing properties

To characterize the firing properties of a neuron, the *rheobase, firing frequency at 700 pA, frequency/current slope (F/I slope)* and the *interspike interval adaptation index (ISI adaptation)* were extracted. The first step current stimulus that elicited at least one AP is referred to as the *rheobase*.The *firing frequency at 700 pA* is the frequency of APs elicited in response to a 700 pA step. The *F/I slope* is the slope of a linear fit through the firing frequencies at increasing step current amplitudes (50-100 pA increments). To calculate the *ISI adaptation*, the first five step currents that elicited at least three APs were used^27^. The *ISI adaptation* was calculated separately for each current injection by dividing the second inter-spike interval (interval between second and third AP) by the first (interval between the first and second AP). The mean value of those five ratios was taken as the final *ISI adaptation* (if less than five values were available, the mean of all available values was calculated).

#### Quality control

To exclude unhealthy neurons and suboptimal recording conditions, we only included neurons for cellular property (Figure S1) and cluster analyses when they met the following criteria:

- Resting membrane potential between −90 mV and −60 mV (without liquid junction potential correction)
- Access resistance < 50 MΩ (24 ± 10 MΩ, mean ± s.d.)
- Laminar depth < 1200 μm

Of 1560 whole-cell patch-clamp recordings, 1096 neurons met these inclusion criteria. For 920 of these neurons, all 15 electrophysiological properties as well as a value for the laminar depth could be extracted and they were used for clustering analyses. For most analyses we focused exclusively on pyramidal neurons. Of the 1560 recorded neurons, 81 were labelled as interneurons by experimenters based on cellular electrophysiological properties or inhibitory synaptic output. For 50 interneurons with successful anatomical staining this labelling was confirmed by the neuron’s morphology (see below). Furthermore, experimenter labelling was confirmed by unsupervised clustering of 920 neurons with complete data, which objectively identified a group of 13 neurons with interneuron characteristics (see below). Excluding the 81 interneurons yielded a pyramidal neuron dataset of 1479 neurons (Figure S1). For cellular property analyses of pyramidal neurons, we used 901 cells, which fulfilled the inclusion criteria listed above and had complete data (Figure S1). For a subset of analyses, we applied looser inclusion criteria and included pyramidal neurons with missing data values for cellular properties (n=163), resting membrane voltage between −60mV and −50mV (n=109) and cells with measured access resistance > 50 MΩ but decent electrophysiological recordings upon visual inspection by experienced patch-clamp electrophysiologists (n=41), yielding a dataset with 1214 pyramidal neurons (Figure S1).

### Synaptic physiology analysis

#### Detection of monosynaptic connections

As described previously^11^, we probed for monosynaptic connections between up to ten simultaneously recorded neurons. In a subset of experiments, we also used the “clean2extend” approach to reuse pipettes that were cleaned in *Alconox* solution to subsequently patch additional neurons^11,60^. During the connectivity recording protocol, all neurons were held close to −60 mV using the automated current injection mode of the MultiClamp Commander. To screen for synaptic connections, we elicited four action potentials in each neuron at 20 Hz with stimulation pulses of 1-4 nA for 1-3 ms in current clamp mode. The minimum stimulation amplitude and duration needed to elicit an action potential was determined prior to connectivity screening. All neurons were stimulated subsequently with an inter-stimulus interval of 1.5 s. That means that, depending on the recording setup, the same neuron was stimulated every 12 or 15 seconds. A monosynaptic connection was identified when correlated postsynaptic potentials could be detected in averaged current clamp traces from 20 to 50 sweeps in any of the other simultaneously recorded neurons. Postsynaptic neurons that did not show a correlated response to presynaptic stimulation were classified as unconnected. All identified monosynaptic connections underwent additional quality control and data curation (see below).

#### Properties of EPSPs

We developed custom MATLAB scripts to automatically extract features of the EPSPs from the raw files. We first computed the EPSP baseline (10 ms interval before presynaptic AP stimulus), EPSP amplitude (peak of moving mean of 0.5 ms sliding window in interval of 0 to 20 ms after maximum upstroke time of presynaptic AP) and the amplitude and peak time of the presynaptic AP in single sweep current clamp traces. We used these sweepwise parameters to automatically exclude sweeps with bad recording quality based on the following exclusion criteria: Postsynaptic baseline more positive than −45 mV or more negative than −80 mV, EPSP amplitude above 20 mV (excludes postsynaptic APs), presynaptic AP amplitude below −50 mV (potentially not triggered AP), presynaptic AP peak time within 0.5 ms of stimulus (avoids stimulus artifact).

After automated sweep exclusion, we computed synaptic parameters of each identified connection based on averaged EPSP traces of the included sweeps (in most cases 30 to 50 sweeps). The EPSP baseline was calculated based on the average of 10 ms before AP stimulus. EPSP amplitude was determined based on the peak of a moving mean (0.5 ms sliding window) in an interval between 0.7 ms (avoid stimulus artifact) to 20 ms after maximum slope time of presynaptic AP relative to the baseline. The same procedure was used to compute the amplitude of the second EPSP that is elicited by the second presynaptic AP. The paired pulse ratio was calculated as the ratio of the second to the first EPSP. The EPSP latency was determined as the time between maximum slope time of the presynaptic AP and the EPSP onset time. The EPSP onset was automatically determined by a series of steps: 1) Find negative peak of postsynaptic trace 0.7 ms after stimulation (avoid stimulation artifact) and before EPSP peak as start of detection interval, 2) use maximum slope time of EPSP as end of detection interval, 3) calculate a moving mean (0.5 ms sliding window) of the derivative of the averaged postsynaptic EPSP (= smoothed slope), 4) identify EPSP onset as the time point when the smoothed slope exceeds three times of the standard deviation during baseline (calculated on derivative in 30 ms interval before next AP). These steps allowed reliable detection of the EPSP onset and thereby synaptic latency which was verified in the manual curation process (see below). Note that synaptic latencies below 0.7 ms could not be discerned due to overlap with stimulation artifacts and were assigned the value of 0.7 ms. The EPSP maximum slope was calculated within the interval between negative peak (see step 1 above) and EPSP peak time. The EPSP halfwidth was calculated as the time between the EPSP trace crossed above and below 50% of its peak amplitude. Note that it was only computed in those connections where the EPSP decayed below 50% of peak within the 50 ms inter-stimulus interval.

#### Curation of synaptic dataset

To verify the automatic parameter extraction and to ensure a high quality of our synaptic dataset, we manually curated every single neuron pair that was classified as being connected during the initial connectivity screening as part of the recordings. Synapse curation was performed by visual inspection of plots generated to display both the raw and averaged traces and the automatically extracted parameters. Curation decisions were made in a consensus process by a group of five highly experienced patch-clamp electrophysiologists (H.P., F.M., H.A., J.G. and Y.P.). Of all initially classified 1466 monosynaptic connections, 450 connections were flagged and underwent manual data curation (the following categories are not mutually exclusive): 47 connections were determined as false-positives and relabelled as ‘not connected’. We curated the synaptic amplitude of 245 connections: 53 connections did not have sufficient recording quality and we removed the amplitude value (e.g. due to depolarized postsynaptic membrane potential or too few sweeps), 92 connections required either manual sweep selection or remeasurement using the respective commercial recording software. We curated the synaptic latency of 94 connections: 65 connections did not have sufficient recording quality and we removed the latency value, 29 connections required manual remeasurement. We curated the synaptic halfwidth of 197 connections: All of these did not have sufficient recording quality to reliably detect halfwidth and we removed the halfwidth value, no manual remeasurement was attempted for this parameter. 46 connections were identified as exhibiting di- or polysynaptic interference on top of the monosynaptic response and were thus only used for the connection probability analysis (Figure S1). The curation decisions and remeasurements were documented in a table and automatically applied to the dataset via custom-written MATLAB scripts. For different analyses of the synaptic dataset, these criteria were specifically combined with cellular and morphological inclusion criteria to ensure strict quality control of each analysis (see respective method section and Figure S1). As a result of the strict quality control and extensive curation procedure, synaptic measurements of the averaged traces were sufficiently reliable, and no model fitting procedure was needed.

### Morphological analysis

#### Cell classification

Out of all 203 recorded clusters, successful anatomical staining was obtained in 169 clusters with a total of 1162 neurons which could be assigned to the electrophysiology dataset (75% of all recorded neurons). These neurons were classified into 1112 pyramidal neurons and 50 interneurons based on their morphology (e.g. dendritic and axonal arborization pattern). The morphologically identified interneurons were excluded from the dataset.

#### Alignment of cells and their laminar positions

During the patch-clamp recordings, relative soma positions were documented using the software of the DIC infrared microscope (Olympus cellSens). These soma location coordinates in the acute slices were then used to align and scale the soma coordinates obtained from the immunohistochemical staining using a custom-written MATLAB script. The distance from the pia vertically to the soma of each neuron was taken as their laminar depth. From 85 neurons no coordinates were obtained during recording; These neurons and their synapses were excluded from depth-based analyses (Figure S1).

#### Detection of slice cutting artifact for connectivity analysis

The slicing procedure of acute brain tissue is necessary for high-resolution patch-clamp recordings. During this process, special care is taken to maintain an optimal cutting angle in order to avoid cutting of dendrites and axons as this leads to underestimation of synaptic connection probability. We optimized the slicing procedure to only cut parallel to the apical dendritic axis (perpendicular to cortical surface). To confirm the parallel cut and the preservation of axons, we estimated the cutting angle of the slices based on the orientation of the apical dendrites relative to the slice surface and assessed the axon of each recorded and stained pyramidal neuron (n = 1112). 25 slices exhibited dendrites that were not parallel to the slice surface with the basal domain and axon pointing towards the slice surface. These slices included 166 pyramidal neurons which were excluded from the connection probability analysis (Figure S1). In the remaining slices, 84 pyramidal neurons had an axon length of less than 100 μm and were also excluded from the connection probability analysis (Figure S1).

### Statistical- and cluster analysis

Statistical analyses and data visualisation were performed using R^61^ and the IDE RStudio^62^. Unless otherwise specified, the ‘tidyverse’ R package^63^ combined with CorelDRAW (Corel Corporation) was utilized to visualize the data. The R packages used for statistical analyses are cited in the respective sections below.

#### Linear regression-, linear mixed-effects- and distribution models

The ‘lm’ function from the ‘stats’ package^61^ was used to fit linear regression models. To test for statistical significance, regression models were compared to intercepts-only models using an F-test. Linear mixed-effects models were fit to the data using the ‘lme4’ package^64^. For analysis of laminar depth dependence of electrophysiological cellular properties (Figure 2, Figure S2), we specified laminar depth as a fixed effect and modeled random intercepts and slopes by human subjects: *‘electrophysiological property ~ laminar depth + (1 + laminar depth | human subject)’*. To obtain the fraction of variance of electrophysiological cellular properties explained by differences between human subjects (Figure S3), we fit random-effects models to our data using the ‘lme4’ package: *‘electrophysiological property ~ (1 | human subject)’*. We calculated the Intraclass Correlation Coefficient (ICC) from these random-effects models using the ‘icc’ function from the ‘performance’ package^65^. The ICC can be interpreted as the fraction of the total variance that is explained by a certain grouping structure in the data, in our case human subjects^39^. The ‘fitdist’ function from the ‘fitdistrplus’ package^66^ was used to fit gaussian and log-normal distributions to the empirical EPSP amplitude distributions of individual human subjects (Figure 3G-H). For each subject the resulting gaussian and log-normal models were then compared using the Akaike Information Criterion (AIC).

#### Correlation- and principal component analysis of cellular properties

The correlation matrix containing Spearman rank correlation coefficients of the 15 electrophysiological properties was calculated using the ‘stats’ package and visualized using the ‘ggcorrplot’ function. The principal component analysis (PCA) of the 15 electrophysiological properties was performed using the ‘prcomp’ function of the ‘stats’ package. Before performing PCA, the data was standardized (Z-scores).

#### Unsupervised hierarchical clustering analysis

We performed unsupervised hierarchical clustering analysis of all neurons that passed quality control and had complete data (n = 920) based on the 15 electrophysiological parameters, using the ‘hclust’ function of the ‘stats’ package. Prior to clustering, the data was standardized (Z-scores). Euclidean distance between data points was calculated and ‘ward.D2’ linkage criterion was used as the agglomeration method^67^. The resulting dendrogram was cut at the level of two groups. One of the two resulting groups contained 13 neurons with clear fast-spiking interneuron characteristics upon manual inspection. The other 907 neurons, which had properties typical for pyramidal neurons, were clustered again using unsupervised hierarchical clustering (Figure 4C, Euclidean distance and ‘ward.D2’ method). The ‘sigclust2’ package was used to test the statistical significance of the hierarchical clustering using a Monte Carlo based approach^43^. This method sequentially tests whether the data at each split corresponds to one (H0) or more multivariate Gaussian distributions (H1) while controlling for the family-wise error rate (significance level was set to alpha = 0.05). To screen for an appropriate number of clusters we used the ‘NbClust’ function from the ‘NbClust’ package^68^ to automatically calculate 26 different indices for determining the number of clusters. The results of these different indices suggested that two, three or four clusters would be most appropriate given our dataset. We further used a resampling method to narrow down the appropriate number of clusters and to address how the clustering result is affected by changes in the dataset (Figure S4)^44^. The approach can be outlined as follows. Step 1: Perform unsupervised hierarchical clustering of the pyramidal neuron dataset into k=n clusters. Step 2: Draw 80% random sample and perform hierarchical clustering of the subsample into k=n clusters. Match the clusters obtained in step 1 & 2 by pairing the medoids based on their Euclidean distance. Step 3: Calculate the fraction of cells in the subsample that were correctly classified into corresponding clusters in step 1 & 2. Repeat step 2 & 3 for 10,000 rounds (Figure S4D&G). This resampling approach was performed for k=3, k=4 and k=5 clusters. We obtained the highest median fraction of correctly reclassified pyramidal neurons when the data was clustered into k=4 clusters (Figure S4G). Taken together, we propose that the cellular parameter space analysed in this study can be best described with four functional groups of pyramidal neurons. To find the most distinguishing cellular properties for pyramidal neuron clusters, we compared each cluster against all other pyramidal neurons for each cellular property and selected the properties yielding the highest the F-values (‘aov’ function from the ‘stats’ package). To validate whether other clustering algorithms yield similar results, we additionally performed k-means clustering (‘stats’ package, Hartigan and Wong method^69^) of the pyramidal neuron dataset into k=4 clusters. We used randomly selected starting points (n=10,000) and allowed 10,000 iterations. The k-means clustering yielded a very similar result compared to hierarchical clustering (Figure S4H). Six of the 907 neurons that were clustered into the pyramidal neuron group in the initial clustering step were labelled as potential interneurons by experimenters and, as a conservative measure, were manually removed after clustering analyses, yielding a pyramidal neuron dataset of 901 neurons (Figure S1). Pyramidal neurons with missing data values for cellular properties (n=163), resting membrane voltage between –60mV and –50mV (n=109) and cells with measured access resistance > 50 MΩ but decent electrophysiological recordings upon visual inspection by experienced patch-clamp electrophysiologists (n=41) were recovered for a subset of analyses (Figure S1). Missing data values were aggregated using the ‘na.aggregate’ function from the ‘zoo’ package^70^. This function replaces missing values by the mean of all available values for a certain parameter. The recovered neurons were then assigned to e-type groups using k-nearest neighbour algorithm from the ‘class’ package^71^ with the hierarchical clustering of the strictly curated dataset acting as the training dataset.

#### UMAP dimensionality reduction

To visualize our 15-dimensional parameter space (15 electrophysiological properties), we used the Uniform Manifold Approximation and Projection (UMAP) technique for dimensionality reduction (‘umap’ package^72,73^). The data was standardized (Z-score) before performing dimensionality reduction. The UMAP figures were plotted using the following hyperparameters: n_neighbors = 100, spread = 0.05, min_dist = 5×10^-5^. To generate patient specific UMAP plots, the coordinates of the UMAP projection of the pooled data were used, but only patient specific data points were visualized.

## Data and software availability

The processed data and the code for analysis and visualization will be made available upon publication (DOI: 10.6084/m9.figshare.21749792).

