## Supplementary Figures 1-5 for "Cellular and Synaptic Diversity of Layer 2-3 Pyramidal Neurons in Human Individuals"

### Supplemental Figures 1-5

■ Applied

| Analysis | Requirements | Figure | N of pyramidal neurons (from n humans) | N of tested synaptic connections (PC-PC) | N of detected PC-PC synapses | Exclusion criteria for cells ① | ② | ③ | ④ | ⑤ | Exclusion criteria for synapses ① | ② | ③ | ④ |
|---|---|---|---|---|---|---|---|---|---|---|---|---|---|---|
| Entire pyramidal neuron dataset | | | 1479 (23) | 9834 | 1419 | | | | | | | | | |
| Cellular property analyses | Reliable cellular properties, no cells >1200 µm laminar depth & available cell coordinates | Fig. 2, 4, Suppl. Fig. 2, 3, 4 | 901 (22) | | | ■ | ■ | ■ | ■ | ■ | | | | |
| Connectivity across humans | No cut axons or unparallel slicing angles (see Methods) & no cells > 1200 µm laminar depth | Fig. 3A | 1262 (22) | 6911 | 1094 | | | | | ■ | ■ | | | |
| Spatial connectivity analysis | No cut axons or unparallel slicing angles (see Methods) & available cell coordinates | Fig. 3C | 1262 (22) | 6586 | 1030 | | | | | ■ | ■ | ■ | | |
| EPSP latency dependence on intersomatic distance | Reliable synaptic properties & available intersomatic distance | Fig. 3D | 1262 (22) | 7732 | 954 | | | | | ■ | | ■ | ■ | ■ |
| Synaptic property analyses | Reliable synaptic properties & available cell coordinates | Fig. 3E-F | 1262 (22) | 8154 | 1010 | | | | | ■ | | | ■ | ■ |
| E-type connectivity | No cut axons or unparallel slicing angles (see Methods) & available e-type label | Fig. 6A-B, Suppl. Fig. 5A-B | 1214 (23) | 5960 | 899 | ‡ | | ■ | | | ■ | | | |
| Synaptic properties of e-types | Reliable synaptic properties & available e-type label | Fig. 6D-H, Suppl. Fig. 5D-J | 1214 (23) | 6966 | 866 (*858, 788, 621) | ‡ | | ■ | | | | | ■ | ■ |

**Exclusion criteria for cells:**

① Access resistance >50 MΩ

② Resting voltage < -90 or > -60 mV

③ Resting voltage < -90 or > -50 mV

④ Missing cellular properties

⑤ Cells w/o coordinates or laminar depth > 1200 µm

**Exclusion criteria for synapses:**

① Presynaptic axon cut or unparallel slicing angle

② Missing intersomatic distance between pre- and postsynaptic neuron

③ Missing synaptic properties

④ Di- or polysynaptic interference

* Synapses with missing value for EPSP latency, -maximum slope and -halfwidth were only excluded for sub-analyses that directly relied on the value (Suppl. Fig. 5E-G)

‡ N=41 neurons with access resistance >50 MΩ but decent ephys. on manual inspection were included in this analysis (see Methods)

**Figure S1: Summary table of the pyramidal neuron dataset and applied exclusion criteria for different analyses.** Table rows correspond to the different analyses performed in this study. The description of the analysis, the requirements needed to perform it and the figure panels that show the results are listed in column one to three, respectively. Columns four to six list the number of neurons, tested pyramidal neuron connections (PC-PC) and found synapses between pyramidal neurons (PC-PC) that remain after the exclusion criteria (shown on the right) were applied to the entire dataset.

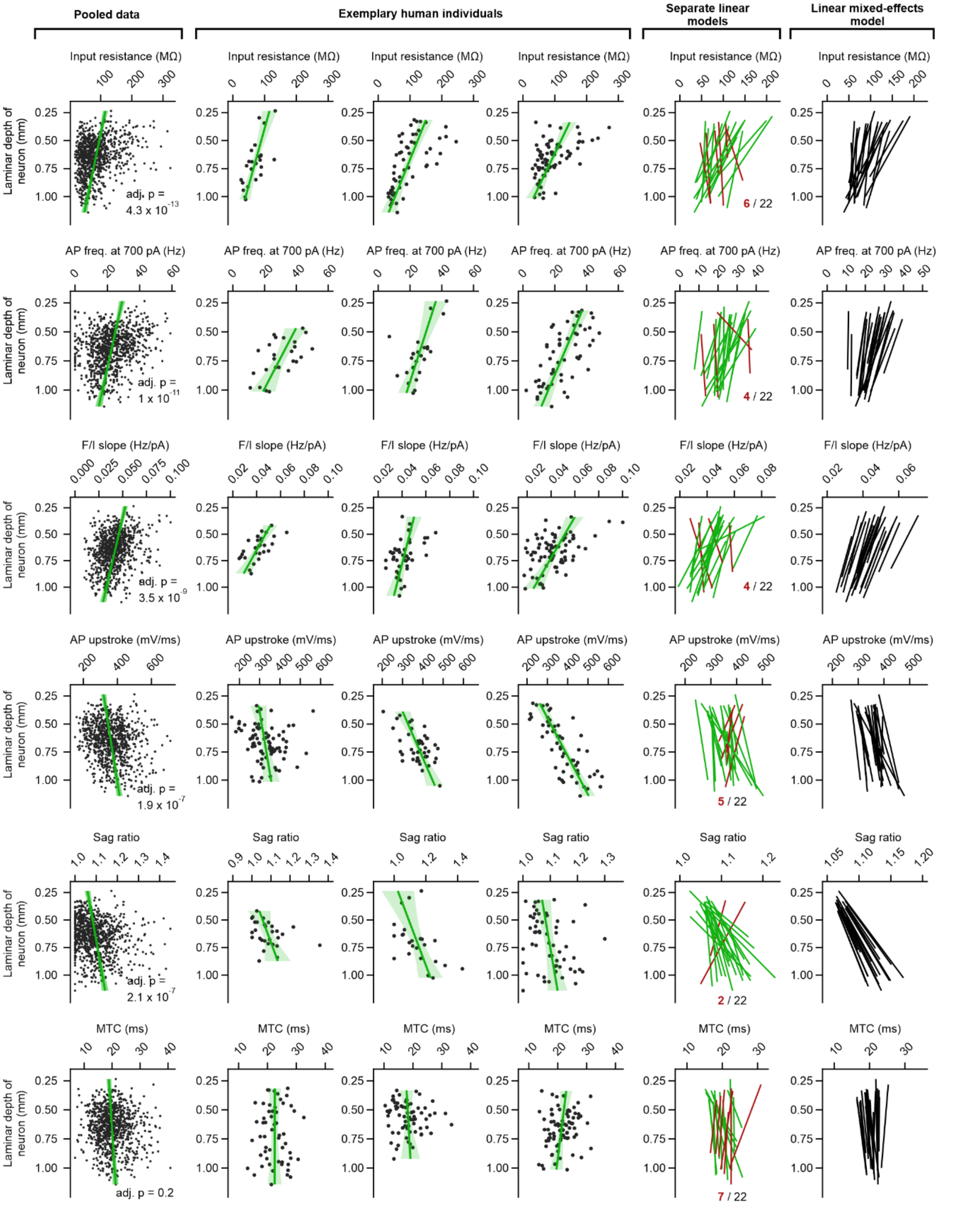


**Figure S2: Laminar depth dependence of intrinsic properties of L2-3 pyramidal neurons.** Each row shows the relationship between laminar depth and one of six selected intrinsic properties. Scatterplots in the first column show pooled data across all human subjects (green line and shaded area correspond to regression line and 95% confidence interval, respectively; F-test, Bonferroni-corrected, adjusted p-values are shown in plots). Columns two to four show relationship between laminar depth and respective intrinsic property in three exemplary human subjects (subjects can differ between rows). The fifth column shows regression lines for each of the 22 subjects. Regression lines with slopes contrary to the pooled data are shown in red. The rightmost column shows fit lines obtained from a linear mixed-effects model with varying intercepts and slopes for each human subject (see Methods).

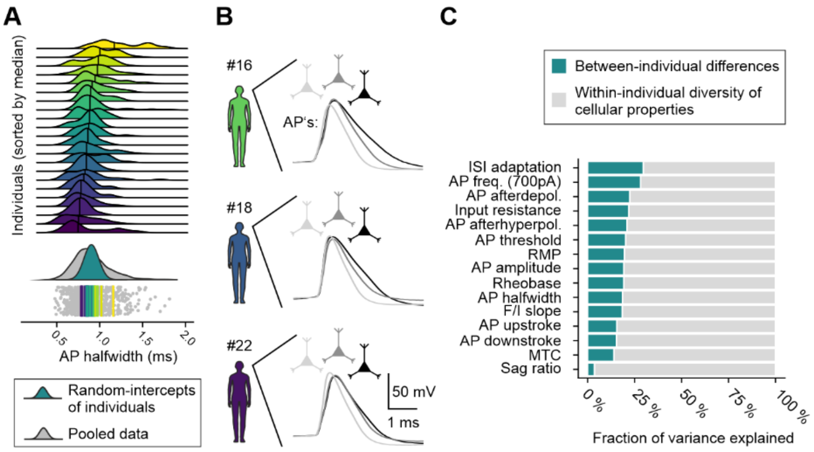


**Figure S3: Between- and within-individual variance**. Action potential (AP) halfwidth (halfwidth of first AP in response to depolarizing currents, see Methods) is shown as an example parameter to demonstrate that within-individual variance of cellular properties is severalfold larger than between-individual variance. (**A**) Top: Smoothed distributions of AP halfwidths are shown for each of the 22 individuals, sorted by descending median (indicated by black vertical lines). Bottom: Jitter-plot and smoothed distribution of pooled data across all individuals (grey). Each dot represents the AP halfwidth of a single pyramidal neuron. Superimposed colored lines and Gaussian distribution represent the random intercept estimates for individuals obtained from a random-effects model (see Methods). (**B**) Schematic visualizing within-individual diversity of AP halfwidths of neurons in three subjects. Note that neurons within the temporal cortex of single humans can have considerably different AP halfwidths. (**C**) Summary bar plot showing fraction of variance explained by differences between individuals (Intraclass correlation coefficient, see Methods) for the 15 electrophysiological parameters in random-effects models.

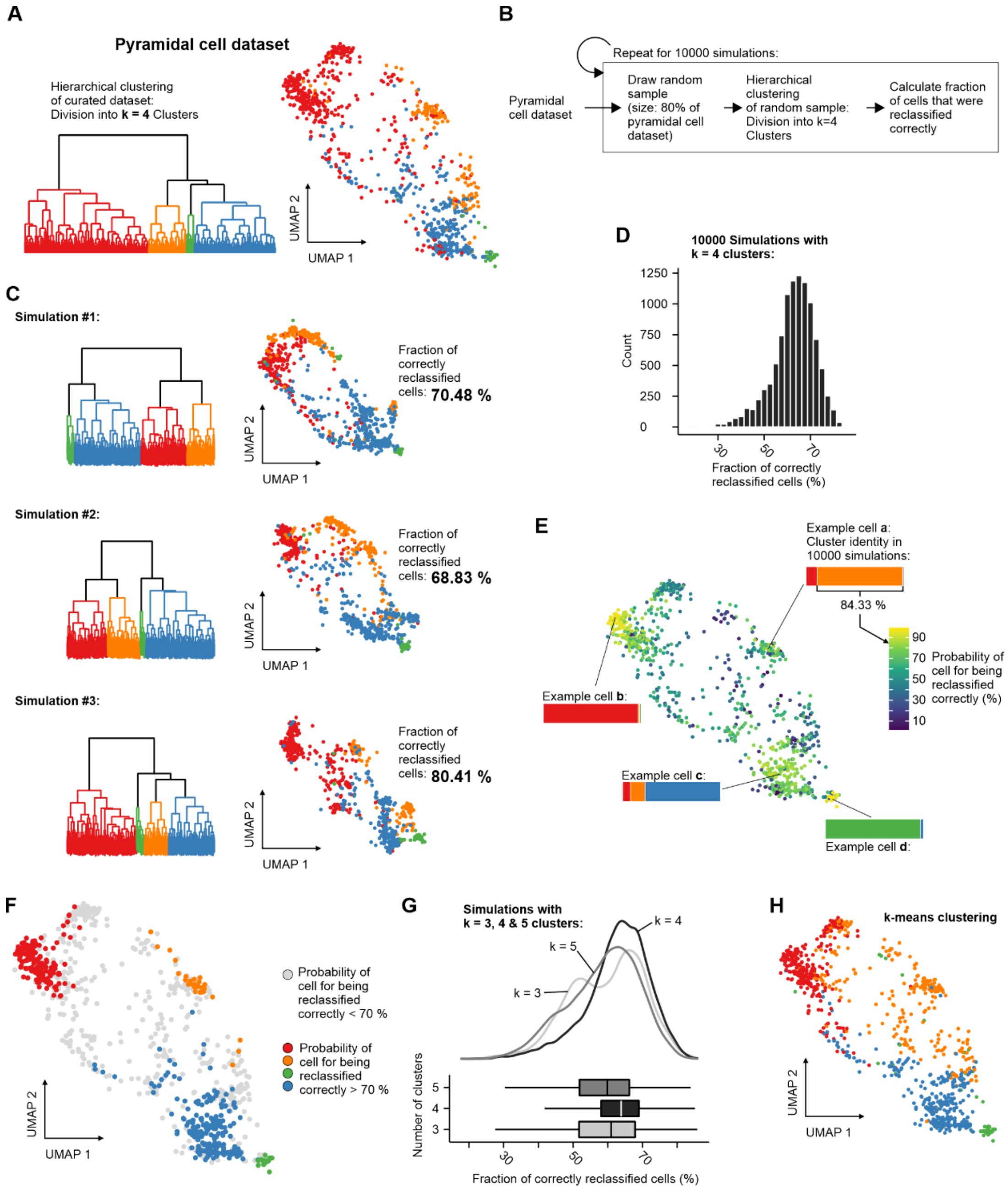


**Figure S4: Resampling analysis of hierarchical clustering.** (**A**) Dendrogram and UMAP plot show the result of unsupervised hierarchical clustering (Euclidean distance, Ward's linkage criterion) of the entire curated pyramidal neuron dataset into four clusters. (**B**) Flow chart depicting the resampling procedure (see Methods). (**C**) Dendrograms and UMAP plots show clustering results of three exemplary simulations with randomly selected 80% subsamples that were clustered into four groups (the fraction of cells that were classified into the same cluster as in the initial clustering shown in A is displayed to the right of the plots). (**D**) Histogram shows the number of correctly reclassified cells in 10,000 simulations. (**E**) UMAP plot color coded by the probability of a cell for being reclassified correctly in 10,000 simulations. (**F**) Same as E, but only cells with a probability > 70% are shown in color. The color corresponds to the cluster that a cell was assigned to in the initial clustering shown in A. (**G**) Smoothed distributions and boxplots show the number of correctly reclassified neurons in 10,000 simulations for k=3, 4 and 5 clusters. Note that simulations in which subsamples were clustered into k=4 groups resulted in a higher median fraction of correctly reclassified cells. (**H**) Clustering result for the entire pyramidal neuron dataset using a k-means clustering algorithm with randomly selected starting points and k=4 clusters (see Methods). Note that the result of the k-means clustering is very similar to the hierarchical clustering shown in A.

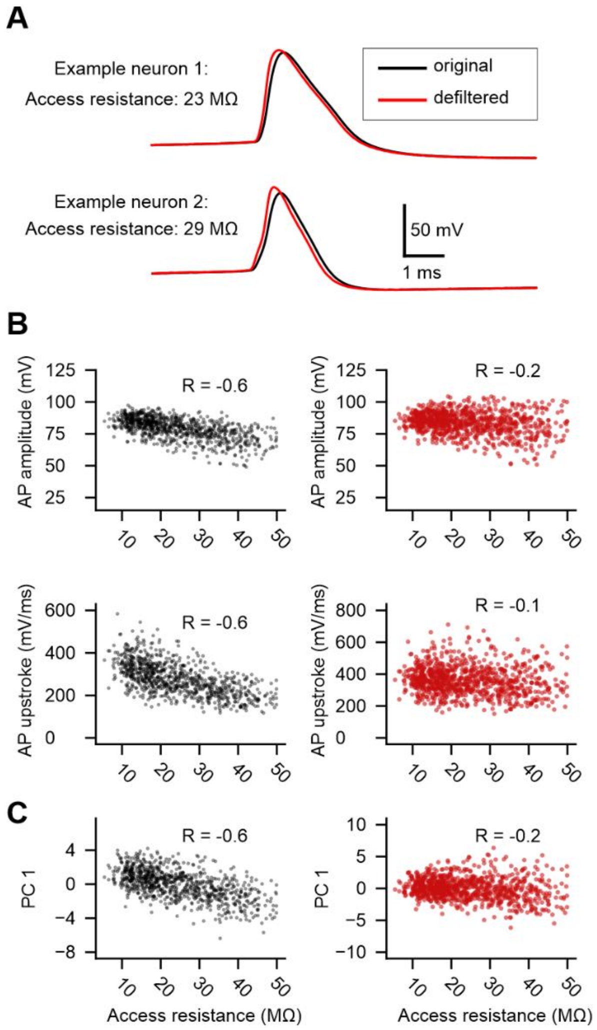


**Figure S5: Defiltering of action potential traces.** (**A**) Action potential (AP) traces of the first AP elicited in response to depolarizing currents of two example pyramidal neurons are shown. The original recorded traces (black) were defiltered using an inverted digital RC-filter (red traces; see Methods). (**B**) Left: AP amplitude and -upstroke extracted from original traces plotted against the access resistance (Pearson correlation coefficient is shown in the plots). Right: Same AP parameters extracted from defiltered AP signals plotted against access resistance. (**C**) First principal component of all seven AP parameters (see Methods) extracted from original (black) and defiltered (red) traces is plotted against access resistance. Note that the correlation is weaker for defiltered signals.
